# Expression of Lysyl Oxidase Family Enzymes During Human Endometrial Decidualization

**DOI:** 10.1101/2025.05.28.656636

**Authors:** Catherine Boniface, Jie Yu, Robert N Taylor, Bronwyn Bryant, John DeWitt, Shanmugasundaram Nallasamy

## Abstract

Lysyl oxidases (LOXs), comprising lysyl oxidase (LOX) and lysyl oxidase like 1-4 (LOXL1-4), constitute a highly conserved enzyme family. Their primary function is the crosslinking of collagen and elastic fibers, thereby modulating the structure and function of the extracellular matrix. Our primary objective was to elucidate the localization of LOXs within human endometrial tissues and to investigate the expression profile of LOXs during *in vitro* human endometrial stromal cell decidualization. Immunohistochemical localization revealed that all LOXs were expressed in human endometrium during both the proliferative and secretory phases of menstrual cycle. All five LOXs were expressed in human endometrial stromal cells cultured *in vitro*. Gene expression levels of *LOX, LOXL1,* and *LOXL2* significantly decreased during endometrial stromal cell decidualization *in vitro*. In contrast, *LOXL3* and *LOXL4* gene expression remained unaffected. Protein levels of LOX, LOXL1, and LOXL3 also showed a significant reduction. Additionally, conditioned media analysis revealed abundant secretion and release of LOX and LOXL1-3 into the extracellular space at all time points examined, with their levels remaining constant. The primary targets of LOXs, collagen and elastic fibers, undergo significant synthesis and processing during endometrial stromal cell decidualization *in vitro*. This observation was supported by the differential gene expression levels of factors involved in their processing and assembly. Collectively, this study demonstrates the expression of LOXs in the human endometrium and their potential role in extracellular matrix reorganization during decidualization.

## INTRODUCTION

Successful establishment of pregnancy involves intricate interactions between the blastocyst and the receptive endometrium [1]. During embryo implantation, the endometrial stromal cells undergo extensive proliferation and differentiation, a process known as decidualization [2]. In humans, decidualization occurs approximately 6 days after ovulation and is characterized by morphological changes in the stromal cells, transformation of uterine glands, vascular remodeling, angiogenesis, and extracellular matrix (ECM) reorganization [3, 4]. ECM, a three-dimensional and dynamic structure, is composed of non-cellular proteins that provide physical support for tissue integrity, facilitate tissue adhesion, mediate signaling, and promote growth. The ECM is unique to each specific tissue, generated embryonically, and allows for specialized and diverse tissue function. Components of ECM are constantly interacting with cells, transmitting signals that control tissue adhesion, migration, differentiation, and apoptosis [5]. Through these complex and tissue-specific interactions, the ECM plays a crucial role in tissue reorganization and homeostasis [6].

ECM, a complex network of macromolecules, comprises collagens, elastin, glycoproteins, proteoglycans, and glycosaminoglycans [5]. Collagen and elastic fibers, the primary fibrous proteins, contribute to tensile strength and tissue resilience, respectively [7, 8]. Lysyl oxidases (LOXs), a family of highly conserved enzymes, are essential for crosslinking and stabilizing collagen and elastic fibers [9–11]. They catalyze the oxidative deamination of lysine and hydroxylysine residues, leading to the formation of highly reactive allysines. These allysines condense, resulting in the formation of mature collagens and elastin crucial for the structural integrity of the ECM [9]. LOXs are divided into lysyl oxidase (LOX) and four lysyl oxidase-like enzymes (LOXL1-4), all of which share a highly conserved catalytic domain at their C-termini. Their diverse functions arise from variations at their N-termini. LOX and LOXL1 contain a pro-peptide sequence involved in their activation upon cleavage. On the other hand, LOXL2-4 contain four tandem repeats of scavenger-receptor-cysteine-rich domains, facilitating protein-protein interactions [10, 11].

ECM reorganization within the decidua provides architectural support and facilitates embryo implantation [3]. Previous studies have demonstrated that different types of collagens were localized in the endometrium throughout the menstrual cycle and in the decidua [12, 13]. Abnormal collagen reorganization and altered lysyl oxidase levels have also been observed in uterine pathologies, including endometriosis and adenomyosis [14–16]. In this study, we examined the expression profile of LOXs in human endometrium and subsequently during *in vitro* human endometrial stromal cell decidualization. We also analyzed the expression of factors involved in the synthesis, processing, and assembly of targets of LOXs, collagen, and elastic fibers.

## MATERIALS AND METHODS

### Tissue Collection

Endometrial curettage samples, collected from premenopausal women in collaboration with the surgical pathology department, were used to localize LOXs. These samples were de-identified and coded. All samples were obtained from women who were free from endometrial polyps, endometrial intraepithelial neoplasia, leiomyomata or malignancies. A postmortem sample of the myometrium collected from a postmenopausal woman consented for participation in research was used as a positive control in immunohistochemical localization of LOXs.

### Immunohistochemistry

Formalin-fixed paraffin-embedded endometrial tissues were sectioned at a thickness of 5 μm. The slides were deparaffinized in xylene, rehydrated through a series of ethanol washes, and rinsed in water. Antigen retrieval was performed by microwave heating for 25 minutes in 0.1 M citrate buffer solution (pH 6.0). The endogenous peroxidase activity was blocked by incubating sections in 0.3% hydrogen peroxide in methanol for 15 minutes at room temperature. The sections were blocked with goat serum and followed by incubating overnight with primary antibodies. The primary antibodies used were LOX antibody (Novus, NB100-2527, 1:100), LOXL1 antibody (ThermoFisher, Scientific PA5-87701, 1:100), LOXL2 antibody (Abcam, ab96233, 1:100), LOXL3 antibody (ThermoFisher, Scientific PA5-48462, 1:100), and LOXL4 antibody (ThermoFisher Scientific, PA5-58436, 1:100). The following day, the tissue sections were incubated with a secondary antibody (Goat anti-rabbit IgG antibody (H+L), biotinylated, ready-to-use) for 30 minutes at room temperature. After washing with TBST, slides were treated with streptavidin for 30 minutes at room temperature. Under the microscope, 3,3’-Diaminobenzidine (DAB) was added to the tissue sections. After optimum staining intensity was observed, slides were immersed in water to stop DAB activity. Slides were counter stained with hematoxylin for 10-15 seconds, rinsed in water for 10 minutes, and immersed in a mixture of NH_4_OH and H_2_O for 30 minutes. After air drying, slides were mounted and imaged using Leica-Aperio VERSA whole slide imager.

### Participant Recruitment and Endometrial Tissue Collection

Healthy, parous women with regular menstrual cycles and no history of hormonal therapy for at least three months prior to surgery were recruited for the study. Endometrial tissue samples were obtained during laparoscopy from participants without evidence of endometriosis or other pelvic pathology. All participants provided written informed consent under protocols approved by the Institutional Review Boards of Wake Forest School of Medicine (No. 00019887) and the Jacobs School of Medicine and Biomedical Sciences, University at Buffalo (No. 00008627). Endometrial biopsies were collected under sterile conditions and immediately transported on ice to the laboratory in Dulbecco’s Modified Eagle Medium/Ham’s F-12 (Catalog No. 10-092CV; CellGro, Manassas, VA) supplemented with 10% fetal bovine serum. Specimens were obtained during the mid-proliferative phase to minimize the effects of endogenous progesterone.

### Preparation of primary human endometrial stromal cells

Primary human endometrial stromal cell (ESC) cultures were derived from mid-proliferative phase biopsies. Cells were sub-cultured at least twice to eliminate potential contamination by macrophages and other leukocytes, following established protocols [17]. ESC cultures generated using this protocol were over 93% pure and retained key phenotypic markers of endometrial stromal cells, including functional estrogen and progesterone receptors, under *in vitro* conditions.

### *In vitro* human endometrial stromal cell decidualization

The stromal cells were cultured in six well plates with DMEM/F-12 medium containing 5%(v/v) fetal bovine serum (Hyclone Laboratories, Logan, UT), 50μg/mL penicillin, and 50μg/mL streptomycin (Invitrogen). After achieving confluency, decidualization was induced as previously described [17, 18]. Briefly, cells were treated with DMEM/F-12 media containing 2% charcoal stripped fetal bovine serum, 50μg/mL penicillin, and 50μg/mL streptomycin and a hormonal cocktail: 10nM E, 1μM P, and 0.5mM 8-bromo-cAMP for 6 days. Media was replaced every 48 hours. Cell lysates and conditioned media were collected at day 0, 2, 4, and 6 after inducing decidualization and stored at −80°C until processing.

### Quantitative PCR

Total RNA was extracted from stromal cell lysates using the RNeasy Mini Kit (Qiagen, 74104) according to the manufacturer’s instructions. cDNA was synthesized using iScript Reverse Transcription Supermix. (Bio-Rad Laboratories). qPCR was performed using SYBR Green master mix and primers specifically designed for the genes of interest. Target gene expression was quantified using the 2^−ΔΔCt^ method and was normalized to the housekeeping gene *RPLP0*. The primers used in this study are listed in Table 1.

**Table 1.**
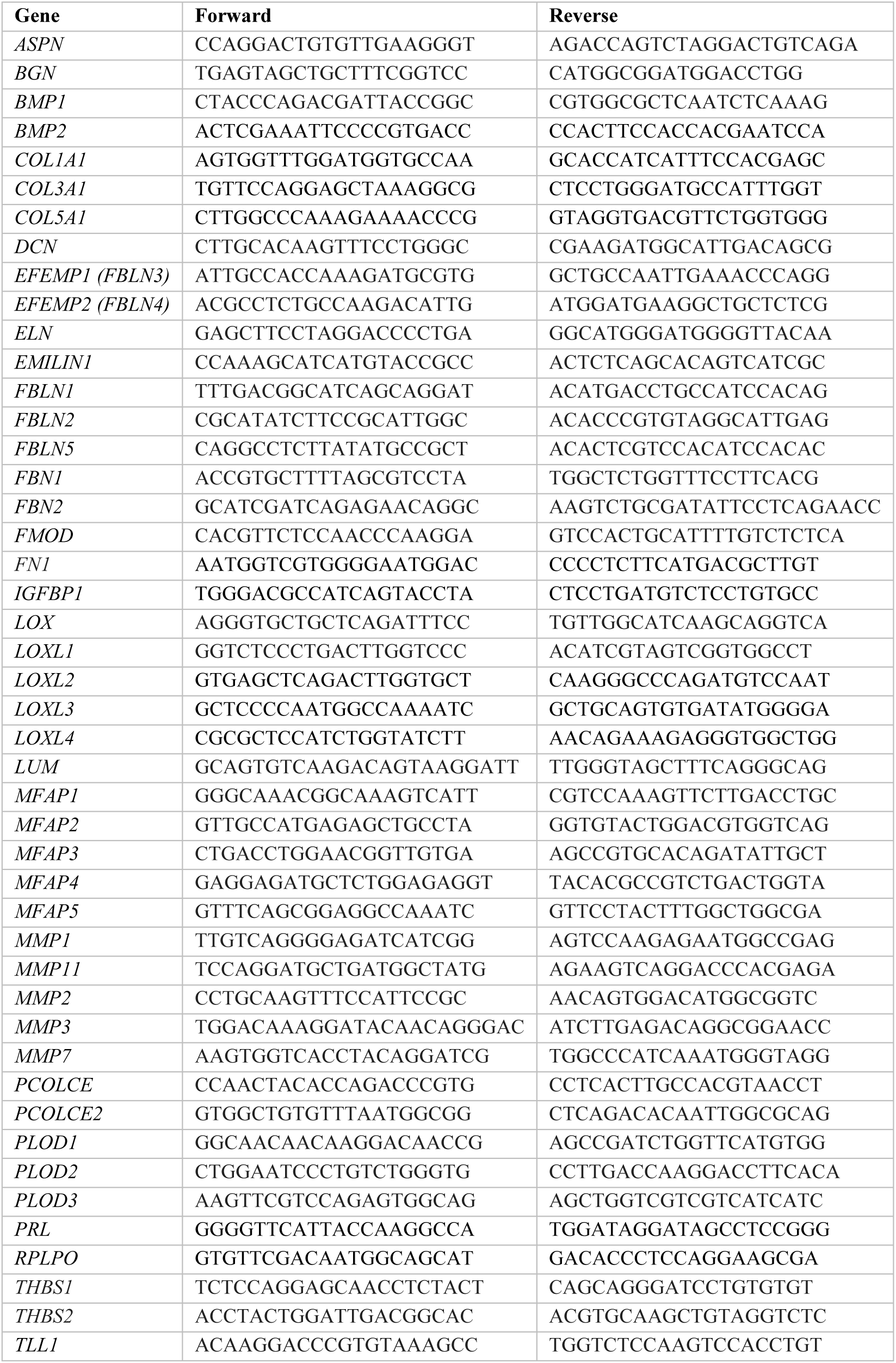
List of primers.

### Western blot

Cells were lysed using RIPA buffer containing 1% protease and phosphatase inhibitors (Thermo Fisher Scientific) for protein analysis. Conditioned media samples were exchanged with equivalent volume of 0.5 M Tris/0.5% SDS, pH 8.0 buffer and concentrated using Vivaspin™ ultrafiltration spin columns (Vivaspin 2, 5 kDa MWCO, # 28932359, Cytiva Life Sciences). Twenty micrograms of protein sample were boiled for 10 minutes at 95°C in 4X Laemmli Sample Buffer (Bio-Rad) containing 5% β-mercaptoethanol. The samples were then loaded into 10% Tris-HCl gel alongside a protein standard (Precision Plus Protein Kaleidoscope, Bio-Rad) and were run at 50V for 30 minutes, followed by 100V for 60 minutes, and then proteins were transferred onto a nitrocellulose membrane (Bio-Rad). Membranes were then blocked using 3% blotting-grade blocker nonfat dry milk in TBST (Bio-Rad) and incubated with a primary antibody in the blocking solution at 4°C overnight.

Primary antibodies used were: COL1A1 (Cell Signaling, 72026, 1:1000), collagen 3 (Proteintech, 22734-1-AP, 1:1000), collagen 5 (Proteintech, 67604-1-Ig), LOX (Abcam, ab174316, 1:1000), LOXL1 (Thermo Fisher Scientific, PA5-87701, 1:500), LOXL2 (Novus, NBP1-32954, 1:500), LOXL3 (Thermo Fisher Scientific, PA5-48462, 1:1000), LOXL4 (Thermo Fisher Scientific, PA5-115520, 1:500), and GAPDH (Cell Signaling, 97166, 1:500). The following day, the membranes were washed and incubated with secondary antibodies conjugated with horseradish peroxidase (HRP). The antibodies used were goat anti-rabbit IgG (H/L) labeled with HRP (Cell Signaling, 7074S, 1:1000) and goat anti-mouse IgG (H/L) labeled with HRP (Cell Signaling, 7076S, 1:1000). Subsequently, the membranes were imaged using an Amersham IQ800 imaging system (Cytiva).

### Statistical analysis

Data were analyzed using GraphPad Prism software. To compare multiple groups, one-way ANOVA followed by Dunnett’s multiple comparisons test was used. Statistical significance was determined at p<0.05.

## RESULTS

### Localization of LOXs in the human cyclical endometrium

To identify LOXs in the human cyclical endometrium, we performed immunohistochemistry on endometrial samples from premenopausal women at both the proliferative and secretory phases. LOXs were consistently expressed in both phases **(Fig. 1).** Additionally, LOXs were localized in both the epithelium and stromal compartments of the endometrium. Among LOXs, LOX, LOXL1, and LOXL2 showed comparable levels in both proliferative and secretory endometrial samples. LOXL3 staining was less intense in the endometrium compared to the other LOXs. LOXL4 exhibited positive staining in both the epithelium and stroma during the proliferative phase but was mostly limited to the stromal compartment during the secretory phase **(Fig. 1).**

**Fig. 1.**
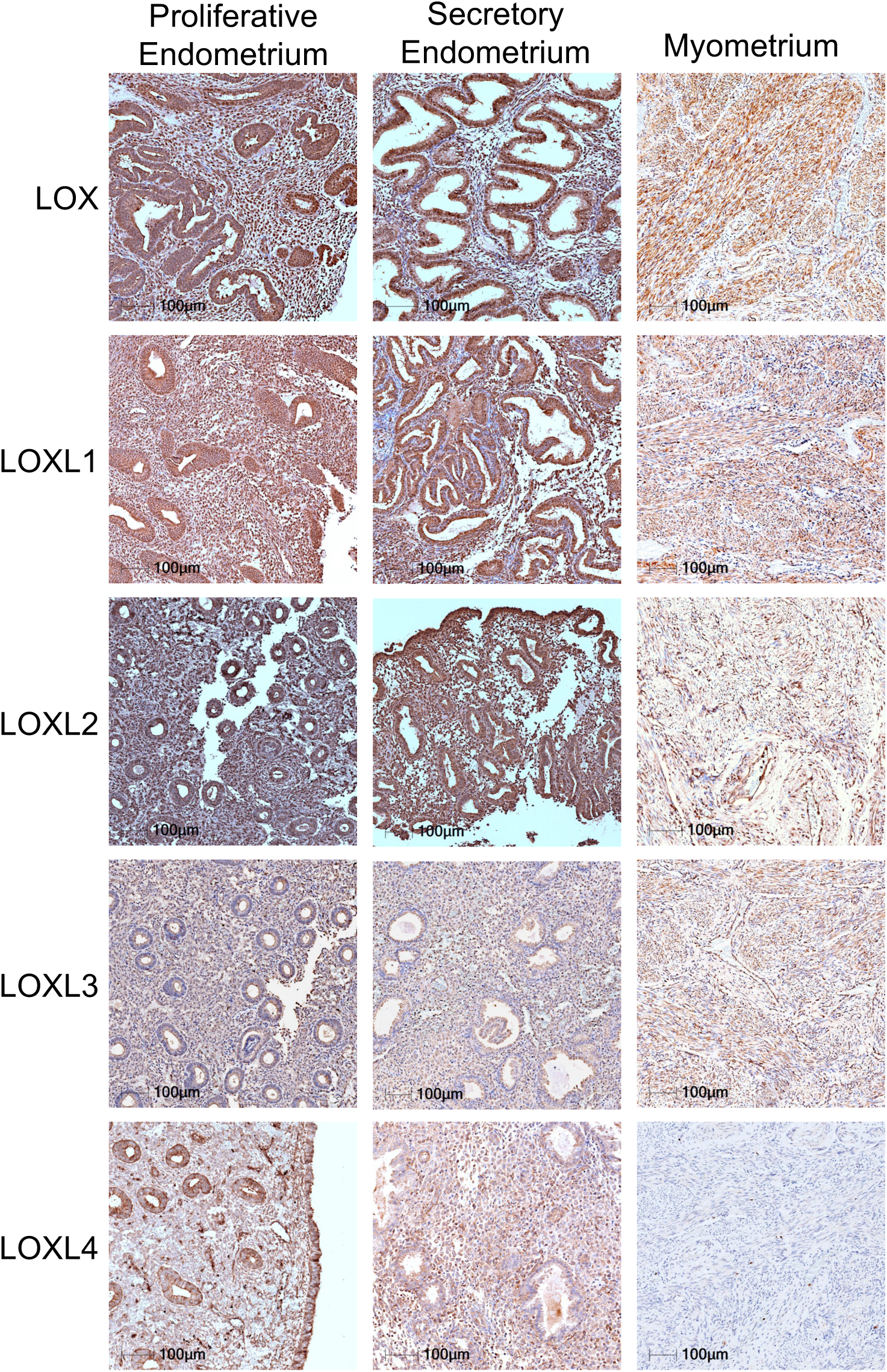
Localization of LOX and LOXL1-4 in human cyclical endometrium. Immunohistochemical staining of LOX and LOXL1-4 in human endometrial tissues collected during the proliferative and secretory phases of the menstrual cycle. Representative images from six independent samples tested. Myometrial tissues served as a positive control. Imaging settings were customized for each individual image to optimize visualization. Scale bar: 100μm.

### Synthesis and secretion of fibrillar collagens during *in vitro* human endometrial stromal cell decidualization

Given the crucial role of LOXs in fibrillar collagen crosslinking, we next analyzed synthesis and secretion of fibrillar collagens 1, 3, and 5 during decidualization. These fibrillar collagens have been localized in the human endometrial tissues [3, 4]. In our study, we examined the synthesis and secretion of these fibrillar collagens during *in vitro* human endometrial stromal cell decidualization. To validate the decidualization process, we used well-established markers of decidualization, including *IGFBP1, PRL,* and *BMP2*, which were significantly induced during decidualization of stromal cells *in vitro* **(Fig. 2A)**. Subsequently, we conducted qPCR analysis of cell lysates to assess the expression of collagen types 1, 3, and 5. Notably, *COL3A1* expression showed a significant increase from day 2, while *COL1A1* and *COL5A1* maintained their levels throughout the process **(Fig. 2B)**. All three collagen protein levels in cell lysates were abundant.

**Fig. 2.**
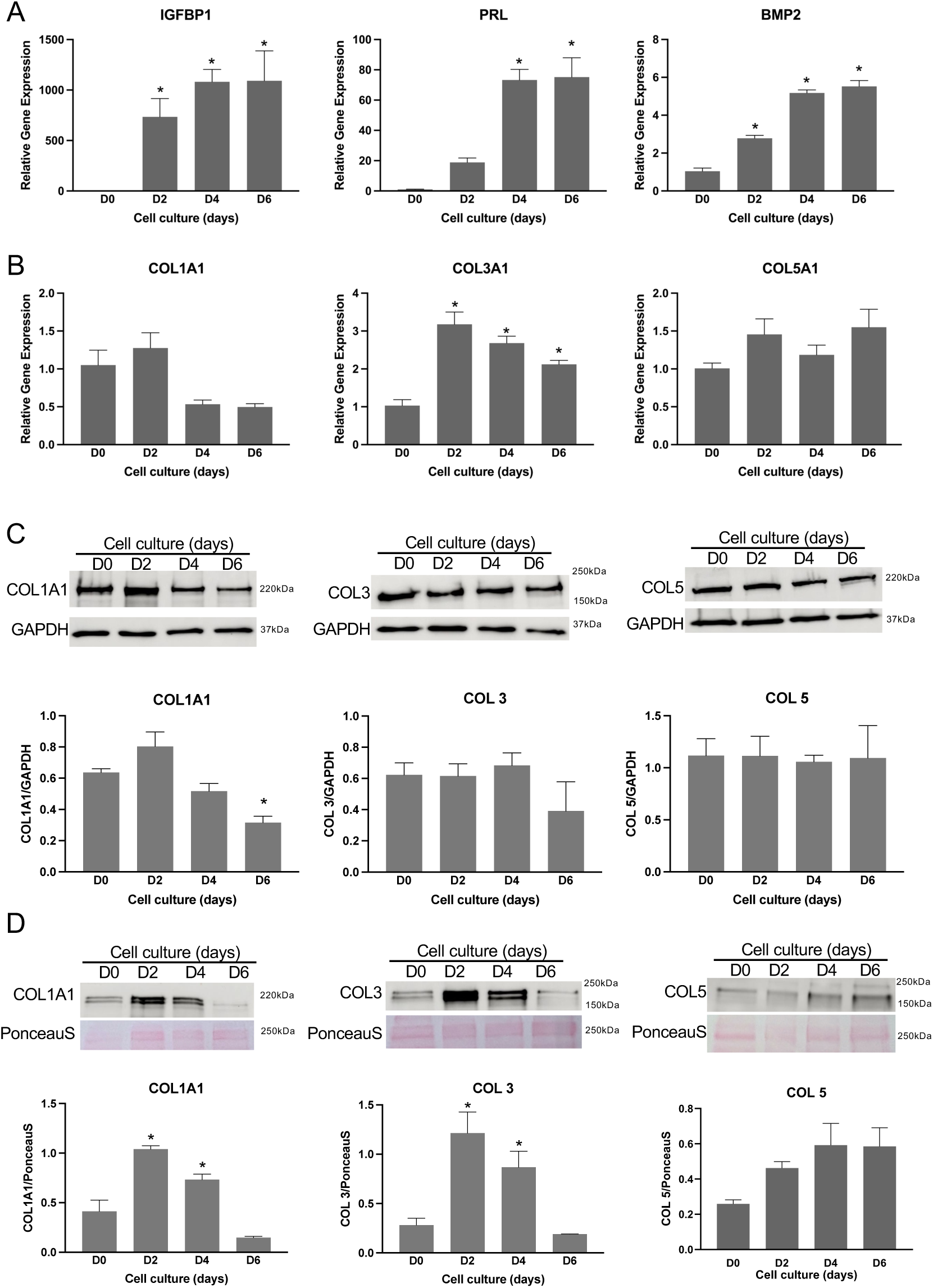
Synthesis and secretion of fibrillar collagens during *in vitro* human endometrial stromal cell decidualization. Primary human endometrial stromal cells were cultured *in vitro* until they reached full confluence. After that, the decidualization was induced with medium containing a hormonal cocktail: 10nM E, 1μM P, and 0.5mM 8-bromo-cAMP for 6 days. The cell lysates were collected at 0, 2, 4 and 6 days to extract total RNA for gene expression analysis. **A.** The decidualization system was validated by analyzing the gene expression of *IGFBP1, PRL,* and *BMP2,* which are known markers of human endometrial stromal cell decidualization. **B.** Gene expression of *COL1A1, COL3A1* and *COL5A1* during *in vitro* human endometrial stromal cell decidualization. The gene expression was normalized to *RPLPO* and compared with day 0 samples (n=4/group, *p<0.05). **C.** Western blot analysis of COL1A1, collagen 3 and collagen 5 protein levels in cell lysates collected at 0, 2, 4 and 6 days. GAPDH was used as a loading control. These are representative images from three to four independent replicates. Quantification of protein levels of COL1A1, collagen 3 and 5 is shown in histogram. **D.** Western blot analysis of COL1A1, collagen 3 and 5 protein levels in the conditioned media collected at 0, 2, 4 and 6 days after the induction of decidualization. These are representative images from three independent replicates. Note: The culture medium was replaced every 48 hours. Ponceau S stain was used to assess the efficiency of protein transfer and visualize the total amount of proteins. Quantification of protein levels of COL1A1, collagen 3 and 5 is shown in histogram (n=3/group, *p<0.05).

In response to decidualization, COL1A1 levels significantly decreased on day 6 compared to day 0, while COL3 and COL5 levels remained constant **(Fig. 2C).** Furthermore, conditioned media analysis revealed a significant increase in COL1A1 and COL3 protein levels day 2 and 4 compared to day 0 **(Fig. 2D).** These findings collectively suggest an enhanced secretion of collagen 1 and 3 as decidualization progresses in human endometrial cells *in vitro*.

The synthesis, processing, and assembly of fibrillar collagen are intricate processes involving various factors, including procollagen proteinases and their enhancers, proteoglycans, matricellular proteins, and fibronectin [7, 19–21]. Beyond their specific function of cross-linking fibrillar collagen, LOXs are known to interact with these factors [22–24]. Therefore, we next investigated the expression of genes encoding crucial factors involved in the synthesis, processing, and assembly of fibrillar collagen. This experiment was aimed to potentially gain insights into the reorganization of fibrillar collagen during decidualization. While the expression of certain proteinases known to facilitate the processing of procollagen into mature collagen, such as bone morphogenetic protein 1 (BMP1), remained stable, the expression of tolloid-like 1 (TLL1) was significantly reduced day 2 **(Fig. 3A).** Interestingly, the expression of enhancers to these proteinases, like procollagen C-endopeptidase enhancer-1 (*PCOLCE*), was significantly induced during decidualization, while the expression of procollagen C-C-endopeptidase enhancer-2 (*PCOLCE2*) remained constant **(Fig. 3A).**

**Fig. 3.**
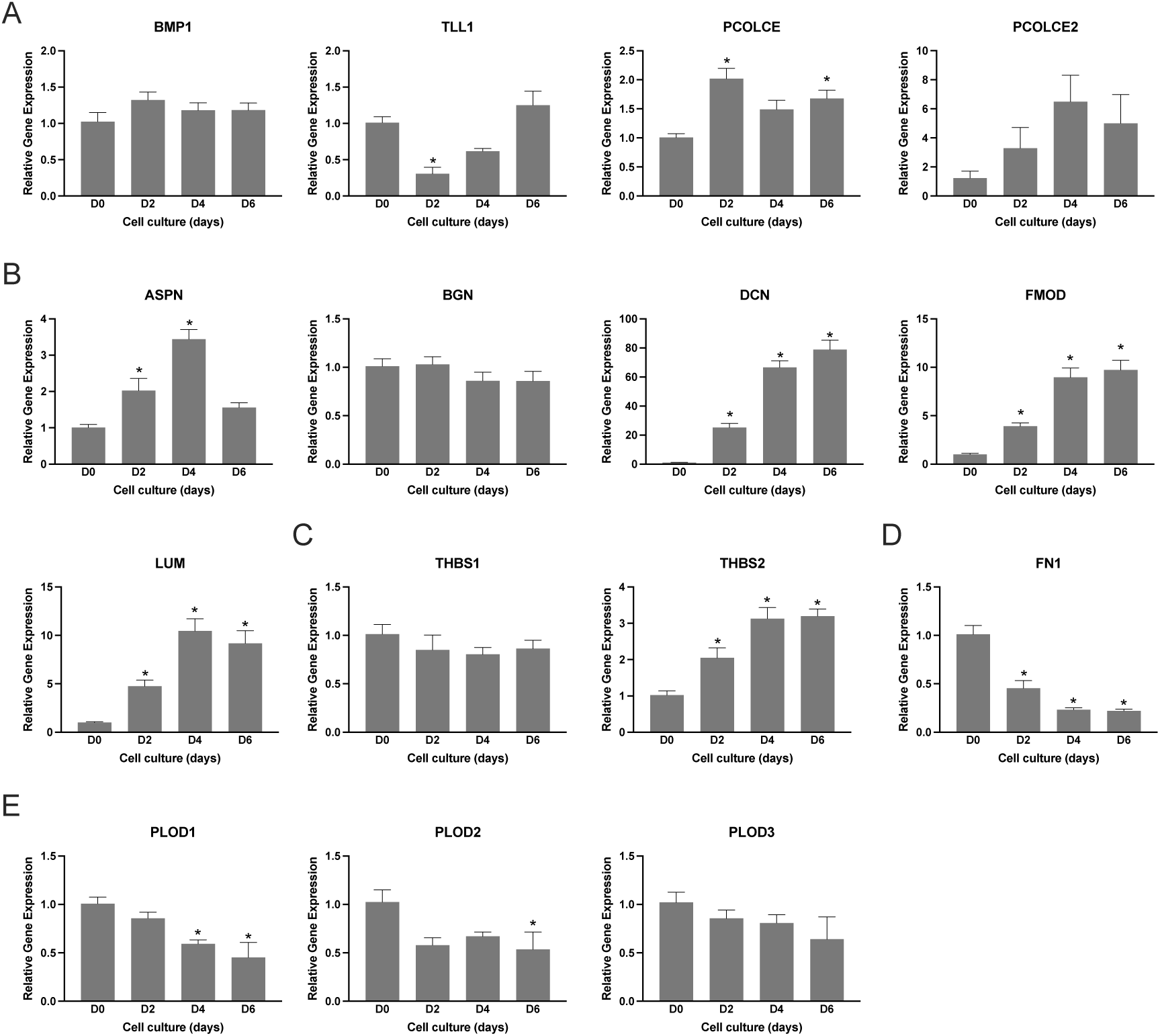
Gene expression of factors involved in the synthesis, processing, and assembly of fibrillar collagen during *in vitro* human endometrial stromal cell decidualization. Primary human endometrial stromal cells were cultured *in vitro* until they reached full confluence. After that, the decidualization was induced with medium containing a hormonal cocktail: 10nM E, 1μM P, and 0.5mM 8-bromo-cAMP for 6 days. The cell lysates were collected at 0, 2, 4 and 6 days to extract total RNA for gene expression analysis. **A.** Gene expression of *BMP1, TLL1, PCOLCE,* and *PCOLCE2*, which encode factors involved in the synthesis and processing of fibrillar collagen, in the human endometrial stromal cells decidualized *in vitro*. **B.** Expression of genes encoding small leucine rich proteoglycans such as *ASPN, BGN, DCN, FMOD,* and *LUM* in the human endometrial stromal cells decidualized *in vitro*. **C.** Expression of genes encoding matricellular proteins such as *THBS1* and *THBS2* in the human endometrial stromal cells decidualized *in vitro*. **D.** Gene expression of *FN1* in the human endometrial stromal cells decidualized *in vitro*. **E.** Expression of genes encoding lysyl hydroxylases, *PLOD1, PLOD2* and *PLOD3* in the human endometrial stromal cells decidualized *in vitro*. The gene expression was normalized to *RPLPO* and compared with day 0 samples (n=4/group, *p<0.05).

We also characterized small leucine-rich proteoglycans (SLRPs), including asporin, biglycan, decorin, fibromodulin, and lumican. These SLRPs play a crucial role in facilitating collagen microfibril growth and assembly into fibrils [20]. SLRPs interact with LOX either directly or indirectly during collagen fibrillogenesis [20, 23]. Many of these proteoglycans showed an increase in expression early during decidualization. Specifically, *ASPN, DCN, FMOD,* and *LUM* were significantly upregulated during decidualization, while *BGN* level remained constant **(Fig. 3B).** Thrombospondin-1 (*THBS1*) and thrombospondin-2 (*THBS2*) are known to modulate cell-matrix interactions and collagen fibrillogenesis [21]. THBS1 in known to interact directly with LOX and collagen [24]. Interestingly, *THBS2* was significantly induced, while THBS1 level remained constant **(Fig. 3C).** Fibronectin (*FN1*) is essential for collagen assembly kinetics [19]. LOXs are essential for fibronectin fibrillogenesis [25]. The expression of *FN1* decreased significantly with the onset of decidualization **(Fig. 3D).**

Lysyl hydroxylases, encoded by *PLOD1-3*, are intracellular enzymes that hydroxylate lysine residues in procollagen, facilitating collagen cross-linking [26]. During endometrial decidualization, *PLOD1* and *PLOD2* levels were significantly reduced, while *PLOD3* remained constant **(Fig. 3E).** Fibrillar collagen reorganization involves both synthesis and degradation, and matrix metalloproteinases (MMPs) are well-known regulators of collagen degradation [27]. Among MMPs, MMP1, MMP2, MMP3, MMP7, and MMP1 are localized in the endometrial tissues [28–30]. Therefore, we analyzed the gene expression of these MMPs during the *in vitro* decidualization of human endometrial stromal cells. All of these MMPs were significantly induced during human endometrial stromal cell decidualization *in vitro*, reaching maximum levels on day 6 **(Fig. 4).** These findings collectively suggest that genes encoding factors influencing collagen synthesis, processing, assembly, and turnover exhibit varying expression patterns. This indicates the dynamic restructuring of fibrillar collagen during endometrial decidualization.

**Fig. 4.**
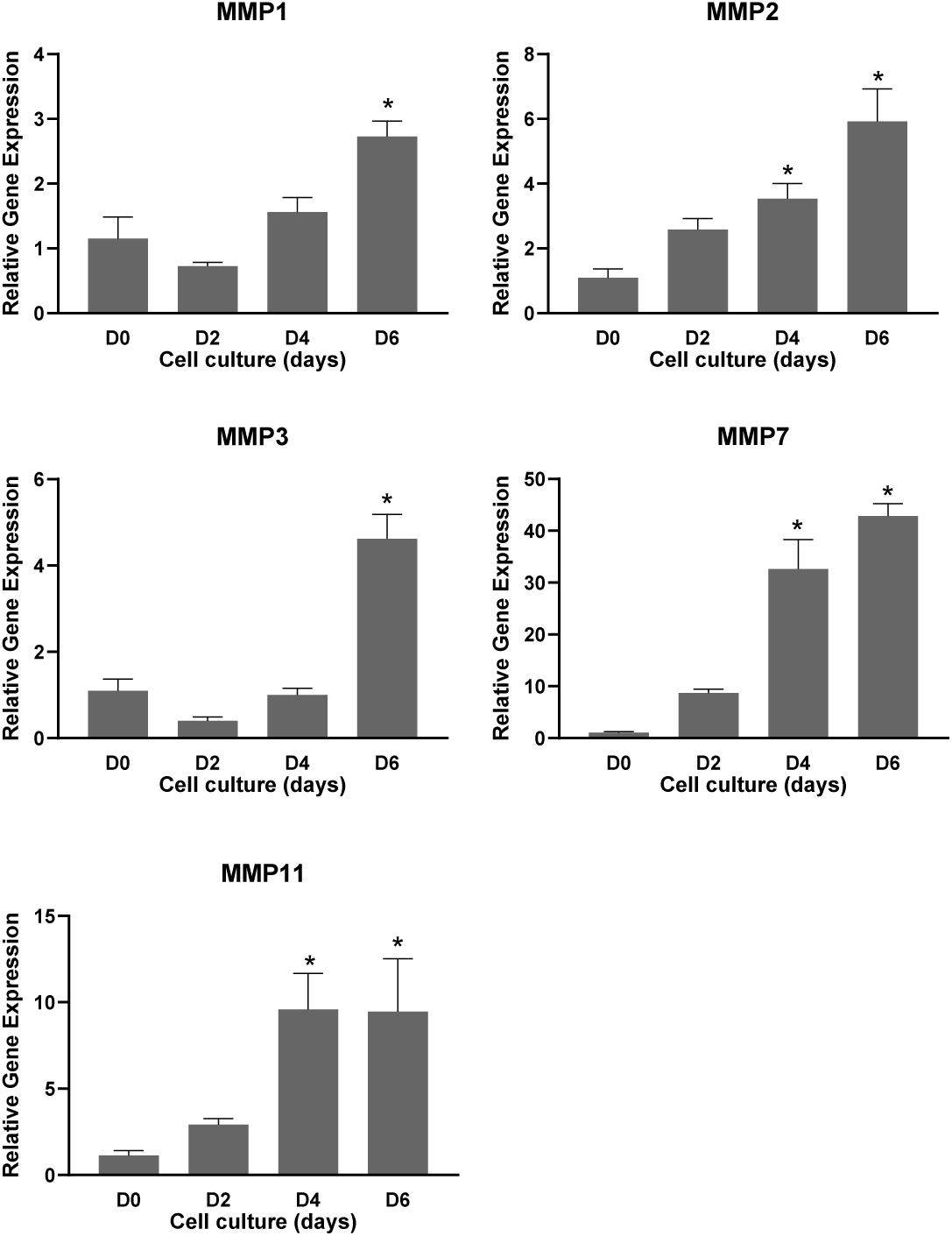
Expression of subset of matrix metalloproteinases during *in vitro* human endometrial stromal cell decidualization. Primary human endometrial stromal cells were cultured *in vitro* until they reached full confluence. After that, the decidualization was induced with medium containing a hormonal cocktail: 10nM E, 1μM P, and 0.5mM 8-bromo-cAMP for 6 days. The cell lysates were collected at 0, 2, 4 and 6 days to extract total RNA for gene expression analysis. Gene expression of *MMP1, MMP2, MMP3, MMP7* and *MMP11* in the human endometrial stromal cells decidualized *in vitro*. The gene expression was normalized to *RPLPO* and compared with day 0 samples (n=4/group, *p<0.05).

### Expression of genes encoding factors involved in the synthesis, processing and assembly of elastic fibers

In addition to fibrillar collagen, LOXs cross-link elastic fibers, contributing to their resilience [31, 32]. Elastic fibers provide elasticity to tissue and are highly concentrated in the vasculature within tissue [8]. Elastin, therefore, is a crucial component of the decidua that undergoes reorganization during embryo invasion to establish the maternal-fetal interface [33]. Similar to fibrillar collagen, the synthesis, processing, and assembly of elastic fibers involve multiple different groups of factors, including fibrillins (FBN1-2), fibulins (FBLN1-5), and microfibril-associated glycoproteins (MFAP1-5) [31]. Therefore, we next analyzed gene expression of elastin and genes encoding factors involved in its synthesis, processing, and assembly in cell lysates obtained from endometrial stromal cell culture. The *ELN* gene, which encodes elastin, was significantly reduced upon induction of decidualization **(Fig. 5A)**. The expression of *FBN1* and *FBN2* genes, which encode structural proteins of microfibrils upon which the elastin is integrated, remained unchanged during decidualization **(Fig. 5B)**. Among the genes which encode fibulins, *FBLN1, FBLN2, FBLN3 (EFEMP1),* and *FBLN5* were significantly induced during endometrial stromal cell decidualization, while *FBLN4* remained unchanged **(Fig. 5C).** Of the genes which encode microfibrillar associated glycoproteins, *MFAP2, MFAP4,* and *MFAP5* were significantly induced during endometrial stromal cell decidualization, while *MFAP3* remained unchanged **(Fig. 5D).** *EMILIN1* (elastin microfibril interfacer 1) remained unchanged during decidualization. **(Fig. 5E).** These findings collectively suggest that genes encoding factors influencing the synthesis, processing, and assembly of elastic fibers exhibit differential expression patterns, indicating that elastic fibers undergo reorganization during decidualization.

**Fig. 5.**
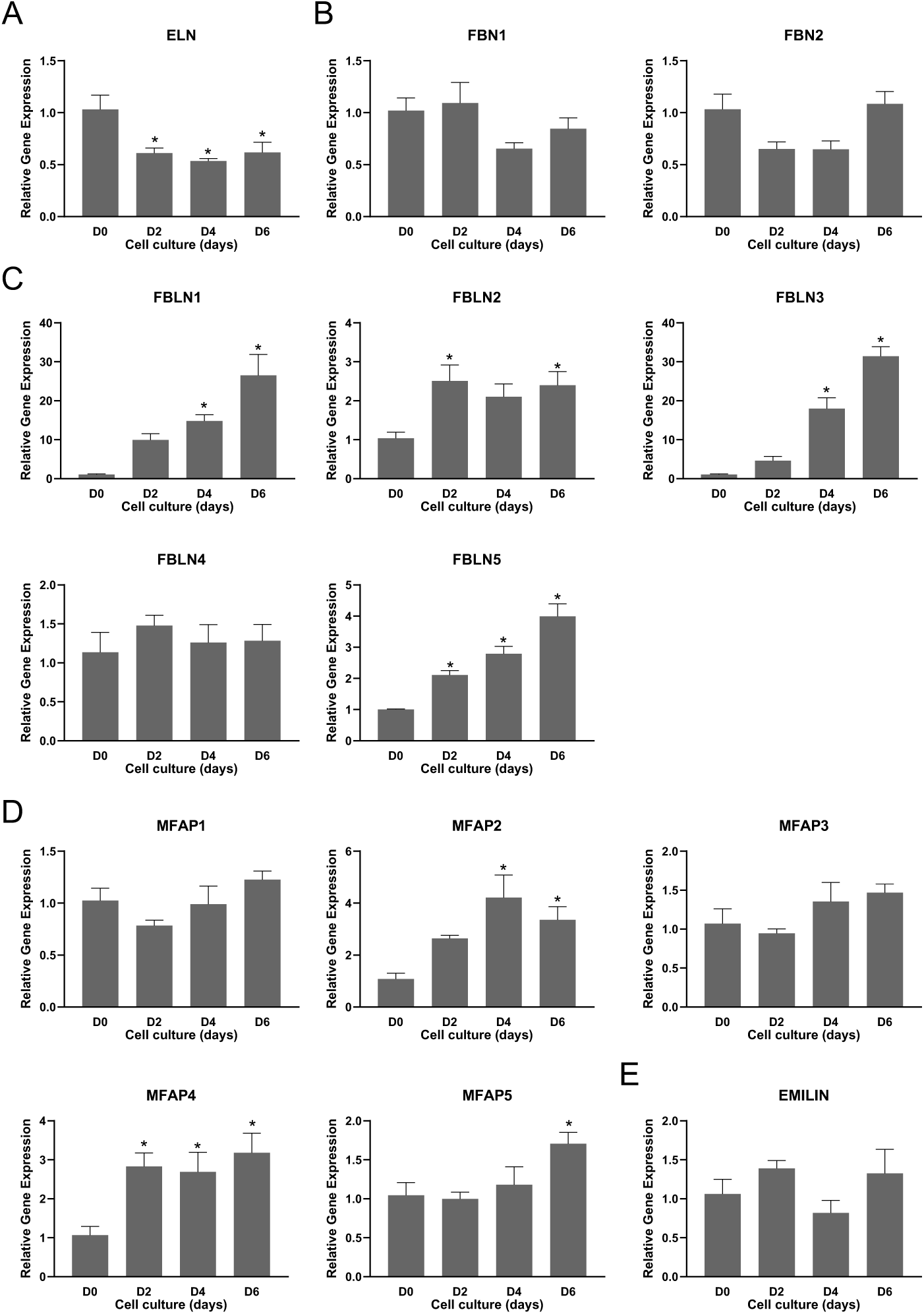
Gene expression of factors involved in the synthesis, processing, and assembly of elastic fibers during *in vitro* human endometrial stromal cell decidualization. Primary human endometrial stromal cells were cultured *in vitro* until they reached full confluence. After that, the decidualization was induced with medium containing a hormonal cocktail: 10nM E, 1μM P, and 0.5mM 8-bromo-cAMP for 6 days. The cell lysates were collected at 0, 2, 4 and 6 days to extract total RNA for gene expression analysis **A.** Expression of *ELN* gene, which encodes tropoelastin monomer, in human endometrial stromal cells decidualized *in vitro.* B. Expression of genes *FBN1* and *FBN2*, which encode microfibrillar proteins, in human endometrial stromal cells decidualized *in vitro* C. Expression of genes *FBLN1-5,* which encode fibulins, in the human endometrial stromal cells decidualized *in vitro*. **D.** Expression of genes *MFAP1-5,* which encode microfibrillar associated proteins, in the human endometrial stromal cells decidualized *in vitro*. **E.** Expression of *EMILIN,* which encode elastin microfibril interfacer-1, in the human endometrial stromal cells decidualized *in vitro*. The gene expression was normalized to *RPLPO* and compared with day 0 samples (n=4/group, *p<0.05).

### Expression of LOXs during *in vitro* human endometrial stromal cell decidualization

To analyze gene expression of LOXs during *in vitro* human stromal cell decidualization, we used quantitative PCR to analyze stromal cell lysates collected from day 0 through 6. All LOXs showed expression in decidualizing cells, but their expression patterns varied **(Fig. 6A).** LOX was significantly reduced by day 2 and 6 compared to day 0. LOXL1 and LOXL2 were significantly reduced upon induction of decidualization, while LOXL3 and LOXL4 maintained their expression throughout the decidualization process **(Fig. 6A).** Utilizing western blot analysis, we could detect all LOXs except LOXL4 in the stromal cell lysates collected during decidualization **(Fig. 6B, C).** Consistent with their gene expression, the protein levels of LOX and LOXL1 were significantly reduced. In contrast, LOXL2 levels remained unchanged, and LOXL3 exhibited significant reduction only on day 6 **(Fig. 6B, C).**

**Fig. 6.**
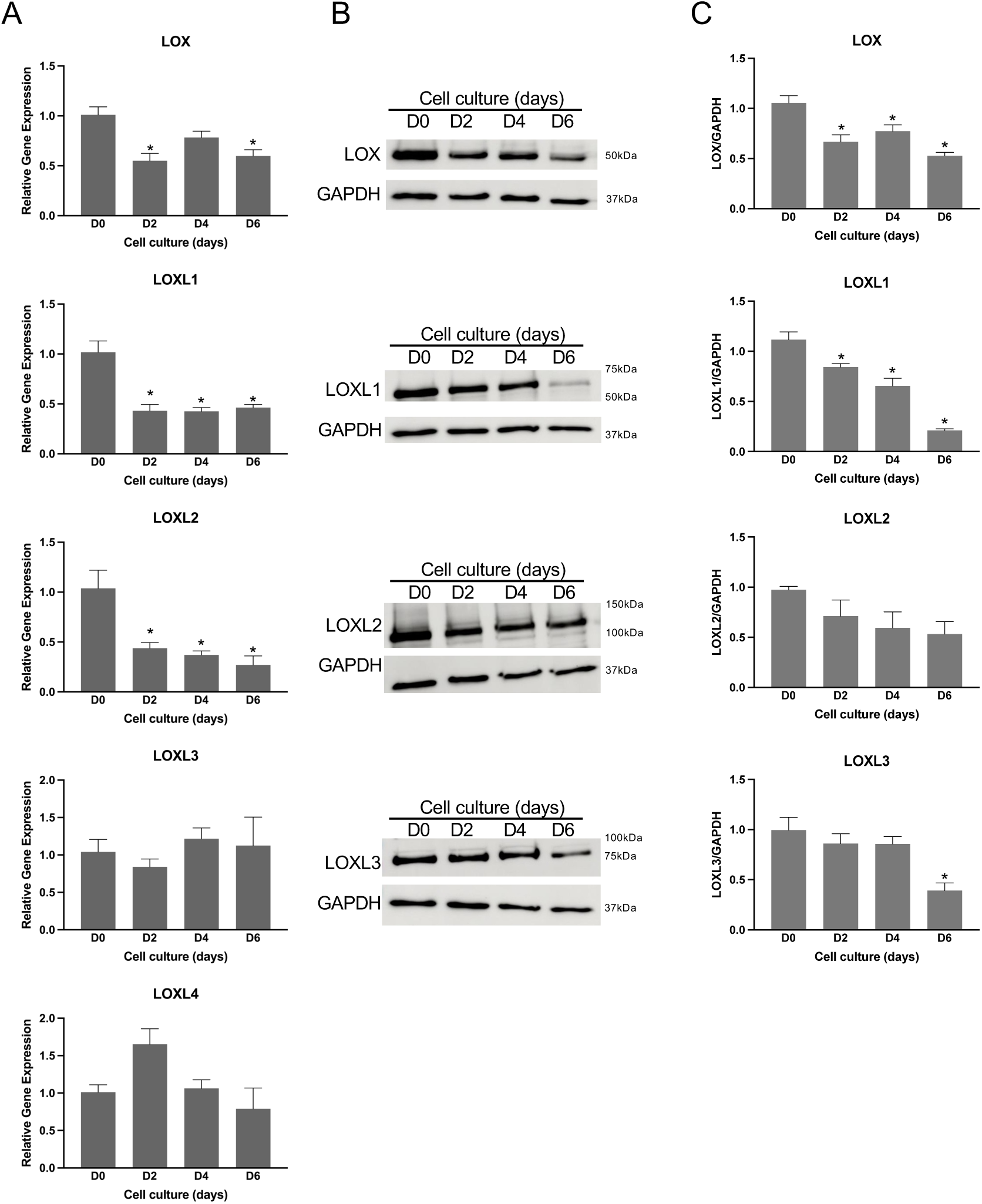
Expression levels of LOX and LOXL1-4 during *in vitro* human endometrial stromal cell decidualization. **A.** Gene expression of *LOX* and *LOXL1-4* in the human endometrial stromal cells decidualized *in vitro*. Primary human endometrial stromal cells were cultured *in vitro* until they reached full confluence. After that, the decidualization was induced with medium containing a hormonal cocktail: 10nM E, 1μM P, and 0.5mM 8-bromo-cAMP for 6 days. The cell lysates were collected at 0, 2, 4 and 6 days to extract total RNA for gene expression analysis. The gene expression was normalized to *RPLPO* and compared with day 0 samples (n=4/group, *p<0.05). **B**. Western blot analysis of LOX and LOXL1-3 protein levels in cell lysates collected at 0, 2, 4 and 6 days. GAPDH was used as a loading control. These are representative images from three to four independent replicates. **C**. Quantification of protein levels of LOX and LOXL1-3 is shown in histogram (n=3/group, *p<0.05).

LOXs are secreted and released into the ECM to undergo processing and mature into functional enzymes. These mature enzymes act on collagen and elastic fibers to mediate crosslinking [11]. Therefore, we next examined the protein levels of LOXs in the conditioned media collected during decidualization using western blot analysis **(Fig. 7A, B).** Similar to cell lysates, we could detect all LOXs except LOXL4 in the conditioned media collected during decidualization. Although LOX and LOXL1-3 were abundantly secreted into the extracellular medium, their levels did not change significantly over the period of decidualization. Collectively, these results suggest that LOXs are synthesized and secreted during *in vitro* human endometrial decidualization and may play a potential role during this process.

**Fig. 7.**
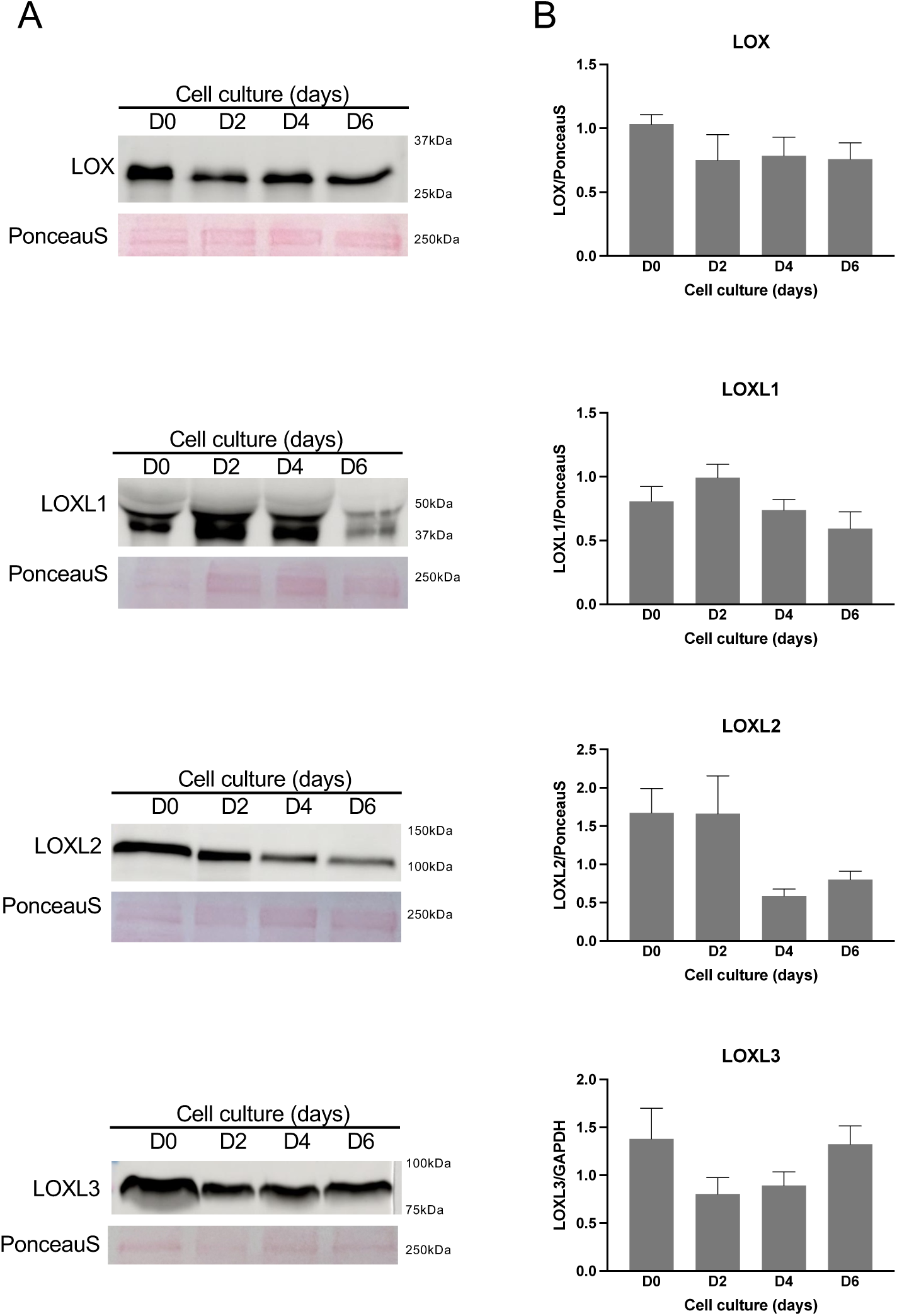
Secretion and release of LOX and LOXL1-3 enzymes into extracellular space during human *in vitro* stromal cell decidualization. **A.** Western blot analysis of LOX and LOXL1-3 protein levels in the conditioned media collected at 0, 2, 4 and 6 days after the induction of decidualization. These are representative images from three independent replicates. Note: The culture medium was replaced every 48 hours. PonceauS stain was used to assess transfer efficiency and to visualize total proteins. **B.** Quantification of protein levels of LOX and LOXL1-3 is shown in histogram (n=3/group, *p<0.05).

## DISCUSSION

During embryo implantation and endometrial decidualization, the endometrial-embryonic interface undergoes extensive reorganization, facilitating the invasion of a growing embryo [1]. ECM reorganization, a crucial process involving multiple factors, transforms this dynamic tissue suitable for embryo invasion [3]. LOX enzymes play a pivotal role in crosslinking and stabilizing collagen and elastic fibers, the primary components of the ECM [10]. Given their central role in ECM reorganization, LOX expression is particularly intriguing during decidualization and implantation. In this study, we investigated the expression profile of LOXs in human cyclical endometrium and human stromal cell decidualization *in vitro*. We also recorded the expression of genes encoding factors involved in the synthesis, processing, and assembly of collagen and elastic fibers, as these are primary targets of LOXs. In cyclical human endometrium, LOXs are expressed in almost all primary cell types, including epithelial and stromal cells, suggesting their potential role in endometrial function. To examine the temporal expression profile of LOXs during endometrial decidualization, we utilized *in vitro* human endometrial decidualization. During this process, most of the LOXs were significantly reduced. These results demonstrate that LOXs are reduced during the ECM reorganization that occurs during decidualization.

The primary functions of LOXs are to cross-link and stabilize collagen and elastic fibers, maintaining tissue architecture and mechanical homeostasis [9, 11]. They also interact with various factors involved in the synthesis, processing, and assembly of these fibers, modifying the structure and function of the ECM in most tissues [22–24]. ECM is uniquely built during embryonic life and undergoes limited reorganization in most adult tissues [6]. In contrast, reproductive tissue ECM undergoes extensive reorganization, including synthesis and turnover, to adapt to their function based on reproductive status [4, 34–36]. The endometrium, for example, undergoes cyclical growth, reorganization, and shedding during the menstrual cycle and extensive reorganization upon embryo implantation to establish the maternal-fetal interface for successful pregnancy [37]. This ECM reorganization is a fundamental event underlying decidualization [3, 4, 37]. While orchestrated reorganization is crucial for successful embryo implantation, aberrant ECM alterations have been linked to pathological conditions such as recurrent miscarriage, endometriosis, adenomyosis, and endometrial aging [4, 14, 38, 39]. Understanding the physiological ECM reorganization may offer clues to understanding disorders caused by abnormal ECM reorganization.

Collagen and elastic fibers, which are crucial for determining tissue integrity and function, are prime targets for LOXs [9–11]. Both collagen and elastin have been found in the human endometrium [3, 4]. Our study demonstrated that fibrillar collagens, particularly collagens 1 and 3, were synthesized significantly during *in vitro* human stromal cell decidualization. This suggests that collagen synthesis contributes to the reorganization that occurs during decidualization. The synthesis, processing, and assembly of collagen and elastic fibers are intricate processes involving multiple factors before they can be assembled to form the ECM scaffold. These fibers undergo extensive posttranscriptional and posttranslational modifications throughout this lengthy process [26, 31]. Analyzing the gene expression of factors involved in these processes provides a valuable estimate of collagen and elastic fiber reorganization. LOXs have been reported to directly interact with these multiple factors, including small leucine-rich proteoglycans such as fibromodulin, fibulins such as fibulin 4 and 5, and matricellular proteins like thrombospondin 1, suggesting their extensive role in determining collagen and elastic fiber organization [22–24]. Our study revealed that various factors involved in these processes were differentially expressed during endometrial decidualization, indicating collagen and elastic fiber reorganization.

LOXs and lysyl hydroxylases play a role in collagen cross-linking [26, 40]. However, they operate at distinct stages of the process and introduce different modifications. Lysyl hydroxylases, intracellular enzymes, specifically hydroxylate lysine residues in procollagen. On the other hand, LOXs enzymes found outside the cell, oxidize lysine residues in both collagen and elastin. These enzymes ultimately form cross-links, which are directly proportional to collagen stiffness and the overall strength of the tissue [26, 40]. During endometrial decidualization, most of the LOXs and lysyl hydroxylases are significantly reduced, suggesting the synthesis of collagen fibers with fewer cross-links. Consequently, the reduction in collagen cross-links indicates that the decidua may reorganize ECM to become softer, facilitating embryo implantation. However, the protein levels of LOXs secreted into the medium were abundant and maintained throughout the decidualization. Therefore, further confirmation through measurements of collagen cross-links is necessary to support this hypothesis.

LOX is dysregulated during uterine pathologies like endometriosis and adenomyosis [15, 16]. Inhibition of the activity of LOXs has been widely considered a feasible therapeutic approach for various conditions, including cancer and fibrosis [10, 41–43]. β-aminoproprionitrile (BAPN), a well-known inhibitor of all LOXs, has been recognized for its irreversible inhibition of all LOXs and has been widely used in experimental research [10, 42, 44]. Additionally, small-molecule selective mechanistic inhibitors of all LOXs, such as PXS-5505 and PXS-4787, have been developed and are currently being tested in preclinical models as therapeutic agents [41, 42]. Therefore, understanding the crucial role of LOXs in reproductive tissues, particularly the endometrium, offers a significant opportunity to develop therapeutic interventions for endometrial disorders.

This investigation of LOX enzymes in human endometrial stromal cell culture has few limitations. In this study, the gene and protein expression results were derived from 2D cell culture. Additionally, studying the spatiotemporal expression of molecular signatures from multiple cell types located at the endometrial-embryonic interface is virtually impossible with *in vitro* models. However, confirming the presence of LOX enzymes in human tissue and documenting their secretion patterns during decidualization serve as crucial starting points for understanding their role in transforming the endometrial extracellular matrix. Future research endeavors will investigate the mechanisms by which these factors regulate endometrial decidualization, thereby providing a more comprehensive understanding of their involvement in endometrial pathophysiology.

## STATEMENTS AND DECLARATIONS

## Author contributions

SN conceptualized and designed the study. CB and SN conducted experiments and analyzed data. CB wrote the first draft of the manuscript. JY and RT provided primary human endometrial stromal cells. BB provided human endometrial tissue samples. JD provided postmortem sample of the myometrium. All authors reviewed, edited and approved the manuscript.

## Acknowledgements

We sincerely acknowledge Dr. Amanda Kallen for her support and guidance during this study. Imaging work was performed at the Microscopy Imaging Center at the University of Vermont (RRID# SCR_018821) and made possible by the College of Medicine Shared Instrumentation Award.

## Ethical approval

All participants provided written informed consent under protocols approved by the Institutional Review Boards of Wake Forest School of Medicine (No. 00019887) and the Jacobs School of Medicine and Biomedical Sciences, University at Buffalo (No. 00008627).

## Data availability

The data underlying this article are available in the article.

